# Leronlimab a humanized anti-CCR5 monoclonal antibody ameliorates hepatic fibrosis in two preclinical fibrosis mouse models

**DOI:** 10.64898/2026.04.17.719186

**Authors:** Melissa Palmer, Taishi Hashiguchi, A. Cyrus Arman, Yuka Shirakata, Neil E. Buss, Jacob P. Lalezari

## Abstract

**Background:** Chemokine receptor type 5 (CCR5) is expressed on hepatic stellate cells (HSCs), which, together with fibroblasts, are major producers of extracellular matrix during liver fibrosis. Leronlimab is a humanized IgG4κ monoclonal antibody that binds to CCR5. The objective of the present study was to evaluate the antifibrotic effects of leronlimab in three independent preclinical studies using two mouse models of liver fibrosis.

**Methods:** In STAM™ (Stelic Animal Model) model 1, leronlimab was administered at doses of 5 or 10 mg/kg/week for 3 weeks. STAM model 2 was conducted as a confirmatory study to validate the antifibrotic effect observed with the 10 mg/kg/week dose in STAM model 1. In a third study, a carbon tetrachloride (CCl₄)-induced liver fibrosis mouse model was used to evaluate leronlimab administered at 10 mg/kg/week for 3 weeks. An isotype-matched control antibody was included in all studies for comparison. Evaluations included liver enzymes and histological assessment of liver fibrosis.

**Results:** In STAM model 1, leronlimab at 10 mg/kg/week significantly reduced fibrosis area compared with the isotype control (p = 0.0005). These findings were confirmed in STAM model 2 (p < 0.0001). Consistent antifibrotic effects were also observed in the CCl₄-induced liver fibrosis model (p = 0.0006).

**Conclusions:** Collectively, these preclinical results demonstrate that CCR5 blockade by leronlimab is associated with a significant reduction of established liver fibrosis in multiple mouse models and support further evaluation of leronlimab as a potential therapeutic option, either as monotherapy or in combination regimens, for chronic liver diseases with fibrosis.

## Introduction

Chronic liver disease of any etiology can progress to fibrosis, ultimately leading to cirrhosis and its associated complications, including hepatocellular carcinoma, liver transplantation, and death. Despite the substantial global burden of fibrotic liver diseases, there are currently no therapies specifically approved to directly target liver fibrosis. Metabolic dysfunction–associated steatohepatitis (MASH) is one of the leading causes of chronic liver disease worldwide [1–6]. Recently, two agents have been approved for patients with MASH and stage 2 or 3 fibrosis: resmetirom, a liver-targeted thyroid hormone receptor β–selective agonist, and semaglutide, a glucagon-like peptide-1 receptor agonist. Phase 3 clinical trials of these agents demonstrated modest improvements in fibrosis compared to placebo, with increased response rates of approximately 12% for resmetirom and 14.4% for semaglutide compared with placebo [7, 8]. Given these limited effects on fibrosis and the fact that neither agent directly targets fibrogenic pathways, there remains a substantial unmet medical need for novel therapeutic strategies, either as monotherapy or in combination, for the treatment of MASH-associated fibrosis and fibrotic liver diseases of other etiologies.

Liver fibrosis is characterized by excessive accumulation of extracellular matrix components [9]. Hepatic stellate cells (HSCs), together with fibroblasts, are the major producers of extracellular matrix and are central to the development of fibrotic scar tissue following chronic liver injury [10].

The recruitment and activation of inflammatory cells at sites of hepatic injury are regulated in large part by chemokines and their receptors, which coordinate immune cell trafficking and signaling within the liver. Chemokine receptor type 5 (CCR5) is expressed on HSCs and is also associated with hepatic macrophage recruitment, inflammatory signaling, and fibrogenesis [11–14]. CCR5 expression has been reported to be upregulated in the livers of patients with MASH [15]. Accumulating evidence implicates CCR5 and its ligands, including CCL5 (regulated on activation, normal T-cell expressed and secreted; RANTES), in the recruitment of monocytes and macrophages, tissue infiltration, and activation of HSCs following liver injury [9–15]. Hepatocytes, Kupffer cells, and infiltrating monocytes/macrophages serve as major sources of transforming growth factor-β (TGF-β), a key profibrotic cytokine that stimulates collagen production by activated HSCs [12]. In addition, CCR2/CCR5 signaling pathways have been implicated in fibrotic processes in other organs, including the kidney, further supporting their role in chronic tissue fibrosis [16–20]. Collectively, these findings highlight CCR5 as a potential therapeutic target in fibrotic liver disease.

Cenicriviroc (CVC), a dual CCR2/CCR5 antagonist, and maraviroc (MVC), a selective CCR5 antagonist, are small-molecule CCR5 inhibitors that have previously been studied in patients with MASLD/MASH. Results from Phase 2 trials demonstrated modest improvements in noninvasive serum and imaging biomarkers [21] and in histologic parameters [22]; however, these findings either did not reach statistical significance [21, 23] or failed to translate into fibrosis regression in phase 3 trials [22–24]. One potential explanation for the limited efficacy of MVC and CVC in MASLD/MASH is incomplete inhibition of chemokine receptor signaling by these small-molecule agents, which may permit residual pathway activity.

Leronlimab (formerly PRO 140) is a humanized IgG4κ monoclonal antibody that binds to extracellular domains of CCR5, including the amino-terminal domain and the second extracellular loop, thereby functionally inhibiting chemokine-mediated CCR5 signaling. Through CCR5 blockade, leronlimab modulates downstream inflammatory and fibrogenic pathways. In a phase 2a partially randomized proof-of-concept clinical study in adults with presumptive MASH without extensive fibrosis [25] leronlimab demonstrated a mild antifibrotic effect in some patients. However, this study was not powered to demonstrate statistical significance on a fibrosis endpoint, histologic samples were not obtained, and the treatment duration of 13 weeks was too short to demonstrate meaningful fibrosis improvement [25]. Thus, evaluation of the antifibrotic effect of leronlimab is needed. In fact, the effects of leronlimab on liver fibrosis have not been systematically evaluated in controlled preclinical models.

The aim of the present study was to investigate the antifibrotic effects of leronlimab in two complementary mouse models of liver fibrosis, including a metabolic injury–driven model and a toxin-induced model, to assess the consistency and robustness of its effects across distinct fibrogenic contexts.

## Materials and methods

### Experimental design

This study was conducted in accordance with the ARRIVE guidelines. Three independent preclinical studies using two mouse models of liver fibrosis were performed to evaluate the antifibrotic effects of leronlimab. These included two proprietary STAM™ (Stelic Animal Model) studies and one carbon tetrachloride (CCl₄)-induced liver fibrosis study. The studies were designed to assess the effects of leronlimab on established liver fibrosis.

### Test substance

Leronlimab, formulation buffer, and an isotype-matched control antibody were provided by CytoDyn Inc. (USA). Details of the test substances are provided in Supplementary Table S1.

### Animals

C57BL/6J mice were obtained from Japan SLC, Inc. For the STAM studies, male offspring derived from 14-day-pregnant females were used, whereas six-week-old female mice were used for the CCl₄-induced fibrosis study. Animals were housed under specific pathogen-free conditions and cared for in accordance with the Japanese Pharmacological Society Guidelines for Animal Use [26–28].

All experimental protocols were reviewed and approved by the Committee on the Ethics of Animal Experiments of SMC Laboratories, Inc. (approval numbers S282 and C044).

### Mouse models

Two independent studies were conducted using SMC Laboratories’ proprietary STAM™ (Stelic Animal Model), which induces insulin resistance and MASH-like liver pathology [25]. In STAM model 1, leronlimab was administered subcutaneously at doses of 5 mg/kg/week or 10 mg/kg/week. STAM model 2 was performed as a confirmatory study to validate the antifibrotic effects observed with the 10 mg/kg/week dose in STAM model 1.

In the STAM models, MASH with fibrosis was induced in male C57BL/6J mice by a single subcutaneous injection of streptozotocin (200 μg; Sigma-Aldrich, USA) two days after birth, followed by feeding with a high-fat diet (57 kcal% fat) from 4 weeks of age. Leronlimab or an isotype-matched control antibody was administered subcutaneously at a volume of 5 mL/kg.

A third study was conducted using a carbon tetrachloride (CCl₄)-induced liver fibrosis model. In this model, female C57BL/6J mice received intraperitoneal injections of CCl₄ twice weekly from Day 0 to Day 35 to induce liver fibrosis, as previously described [29, 30]. Leronlimab was administered at 10 mg/kg/week. CCl₄ is a well-established hepatotoxin that induces hepatocellular injury, inflammatory responses, and progressive liver fibrosis, and is widely used to evaluate antifibrotic interventions.

In all studies, animals were randomized into treatment groups based on body weight to ensure balanced baseline characteristics. For the STAM models, stratification criteria were defined based on historical background data maintained by SMC Laboratories, Inc., whereas for the CCl₄ model stratification was based on baseline body weight measured at study initiation.

During the conduct of the experiments, personnel involved in animal husbandry and dosing were aware of group allocation as required for study execution. Histopathological evaluations were performed by investigators blinded to treatment allocation. Data analyses were conducted using coded datasets, and group identities were revealed only after completion of statistical analyses.

Animals were monitored daily for general health and clinical signs, and body weight was recorded throughout the study. Mice were sacrificed at 12 weeks of age in the STAM studies and on Day 35 in the CCl₄ study by exsanguination under isoflurane anesthesia.

Liver tissues were fixed, paraffin-embedded, and sectioned for histological analyses. Fibrosis was assessed by Sirius red staining, and fibrosis area was quantified using image analysis software (ImageJ, National Institutes of Health, USA) in five fields per section and expressed as a percentage of the total area. Hematoxylin and eosin (H&E) staining was performed for histopathological evaluation. In the STAM studies, NAFLD activity score (NAS) was assessed according to the criteria of Kleiner et al. [31], as summarized in Supplementary Table S2.

All animals allocated to each experimental group were included in the analyses, and no animals or data points were excluded unless explicitly stated.

### Statistics

Statistical analyses were performed using GraphPad Prism version 6 (GraphPad Software, USA). Comparisons between two groups were conducted using unpaired, two-tailed Student’s t-tests. For analyses involving more than two groups, one-way analysis of variance (ANOVA) was applied, followed by Bonferroni post hoc tests where appropriate. Data are presented as mean ± standard deviation (SD). Statistical significance was defined as p < 0.05 (two-sided).

Individual animal data for key endpoints, including Sirius red–positive fibrosis area, NAFLD activity score (NAS), and serum biochemical parameters, are provided in Supplementary Tables S3–S5 to ensure transparency and reproducibility. Effect sizes and corresponding 95% confidence intervals are reported in Supplementary Tables S6–S8.

Formal statistical tests to assess distributional assumptions were not performed; however, data were reviewed for extreme outliers, and the applied parametric tests are consistent with standard practice in comparable preclinical fibrosis studies. This approach is acknowledged as a methodological limitation.

### STAM model 1, STAM model 2, and CCl₄-induced fibrosis trial designs

STAM model 1 consisted of four experimental groups with eight mice per group, and STAM model 2 consisted of three experimental groups with twelve mice per group (Supplementary Fig. S1a and S1b). Sample sizes were determined based on prior experience and background data from studies using the STAM model, which indicated that these group sizes were sufficient to detect consistent differences in histological and biochemical endpoints while minimizing animal use.

In both STAM studies, mice were allocated to the following groups: a normal control group without treatment, an isotype-matched IgG4 control antibody group (10 mg/kg/week), and leronlimab treatment groups. In STAM model 1, leronlimab was administered at 5 mg/kg/week or 10 mg/kg/week, whereas STAM model 2 included only the 10 mg/kg/week leronlimab dose as a confirmatory study. All antibodies were administered subcutaneously once weekly from 9 to 12 weeks of age.

Endpoints evaluated in the STAM studies included liver-to-body weight ratio, plasma alanine aminotransferase (ALT), aspartate aminotransferase (AST), glucose, and triglyceride levels, as well as histological assessment of NAFLD activity score (NAS) and fibrosis area.

The CCl₄-induced liver fibrosis study consisted of three groups with twelve mice per group (Supplementary Fig. S1c). Control animals received intraperitoneal injections of mineral oil twice weekly throughout the study. Fibrosis was induced by intraperitoneal administration of 5% CCl₄ in mineral oil (100 μL) twice weekly. A treatment group received leronlimab at 10 mg/kg/week administered subcutaneously on Days 14, 21, and 28 in addition to CCl₄ administration.

In the CCl₄ study, endpoints included liver-to-body weight ratio, plasma ALT levels, hepatic hydroxyproline content, and histological quantification of fibrosis area.

## Results

### STAM model 1

As shown in Table 1 and 2, the isotype control group exhibited a significant increase in mean liver-to-body weight ratio compared with the normal group. No significant differences in liver-to-body weight ratio were observed between the isotype control group and the leronlimab-treated groups. One mouse in the isotype control group was found dead prior to Day 21 (day of sacrifice); no additional deaths or premature euthanasia occurred.

**Table 1.** STAM model 1: between group differences in parameters.

| **Comparison** | **Bonferroni Multiple**<br>**Comparison test (p-value)** | **Student's t-Test (p-**<br>**value)** |
| --- | --- | --- |
| **Liver-to-body weight ratio** |  |  |
| Normal vs isotype control | <0.0001 | <0.0001 |
| Leronlimab low vs isotype control | >0.9999 | 0.3085 |
| Leronlimab high vs isotype control | >0.0999 | 0.2777 |
| **Whole blood glucose** |  |  |
| Normal vs isotype control | <0.0001 | <0.0001 |
| Leronlimab low vs isotype control | >0.9999 | 0.2806 |
| Leronlimab high vs isotype control | >0.0999 | 0.1513 |
| **Plasma ALT** |  |  |
| Normal vs isotype control | 0.0374 | 0.0366 |
| Leronlimab low vs isotype control | 0.3082 | 0.1245 |
| Leronlimab high vs isotype control | 0.7751 | 0.2139 |
| **Plasma AST** |  |  |
| Normal vs isotype control | 0.0117 | 0.0107 |
| Leronlimab low vs isotype control | 0.2997 | 0.1124 |
| Leronlimab high vs isotype control | 0.3492 | 0.1141 |
| **Plasma triglyceride** |  |  |
| Normal vs isotype control | 0.0417 | 0.0045 |
| Leronlimab low vs isotype control | 0.2242 | 0.0271 |
| Leronlimab high vs isotype control | >0.9999 | 0.3338 |
| Leronlimab low = 5 mg/kg/week; leronlimab high = 10 mg/kg/week. |  |  |

**Table 2.** STAM model 1: Mean ± SD of each parameter.

| **Group** | **n** | **Liver-to-body**<br>**weight ratio**<br>**(%)** | **Whole blood**<br>**glucose**<br>**(mg/dL)** | **Plasma ALT**<br>**(U/L)** | **Plasma AST**<br>**(U/L)** | **Plasma**<br>**triglyceride**<br>**(mg/dL)** |
| --- | --- | --- | --- | --- | --- | --- |
| **Normal** | **8** | **4.6 ± 0.2** | **194 ± 25** | **19 ± 2** | **45 ± 13** | **115 ± 39** |
| **Isotype control** | **7** | **8.4 ± 1.4** | **600 ± 107** | **44 ± 36** | **113 ± 72** | **1937 ± 1692** |
| **Leronlimab low** | **8** | **8.1 ± 0.9** | **642 ± 160** | **29 ± 7** | **76 ± 35** | **653 ± 312** |
| **Leronlimab high** | **8** | **8.0 ± 1.2** | **548 ± 80** | **34 ± 8** | **78 ± 28** | **1501 ± 2087** |

Compared with the normal group, plasma liver enzymes, whole blood glucose, and plasma triglyceride (TG) levels were significantly increased in the isotype control group, confirming successful disease induction (Table 1, 2). No significant differences were observed between the isotype control and leronlimab-treated groups for these parameters.

To evaluate disease activity, hematoxylin and eosin (H&E) staining was performed and the NAFLD activity score (NAS) was calculated. As shown in Table 3, the NAS was significantly increased in the isotype control group compared with the normal group. Both leronlimab-treated groups (5 mg/kg/week and 10 mg/kg/week) showed a significant reduction in total NAS compared with the isotype control group. Although each individual NAS component (steatosis, inflammation, and ballooning) showed numerical reductions in the leronlimab-treated groups, none reached statistical significance individually; the reduction in total NAS reflected the combined effect across components.

**Table 3.**
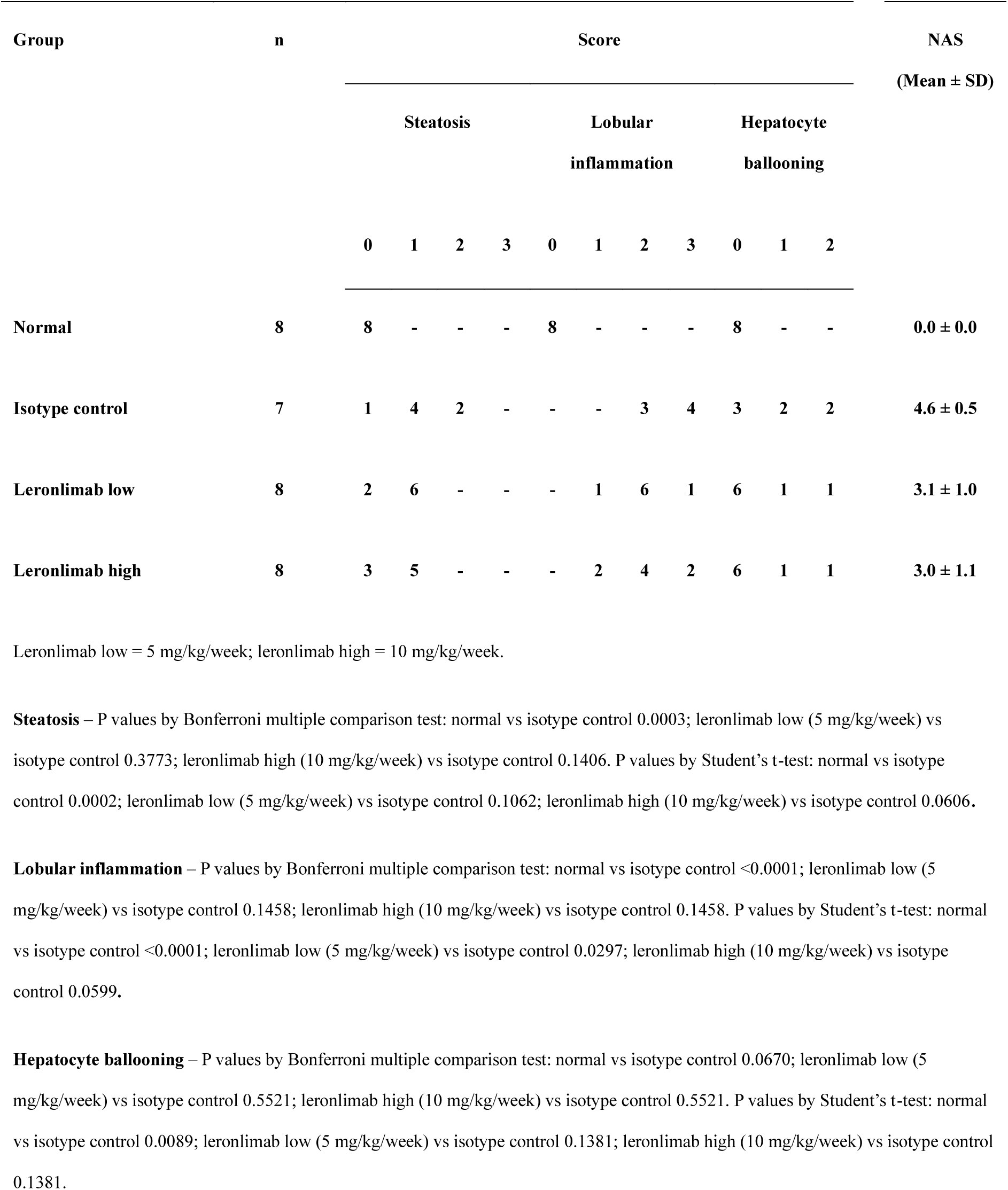
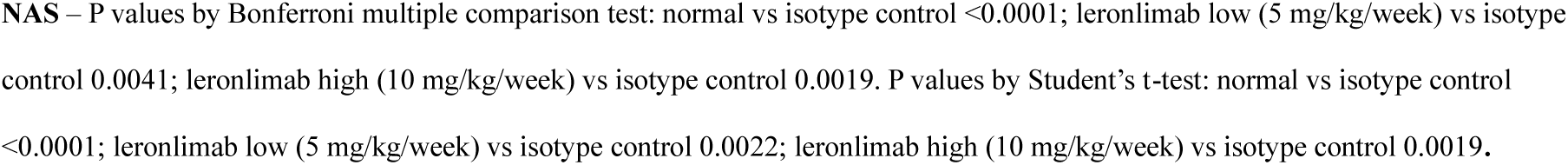
NAFLD activity score STAM model 1.

Histological assessment of fibrosis by Sirius red staining demonstrated a significant reduction in fibrosis area in the leronlimab 10 mg/kg/week group compared with the isotype control group after 3 weeks of treatment, whereas no significant effect was observed with the 5 mg/kg/week dose (Fig. 1a). Representative Sirius red–stained liver sections are shown in Fig. 1b–c. Based on the lack of significant antifibrotic effect at 5 mg/kg/week, this dose was not evaluated in the confirmatory study (STAM model 2).

**Fig. 1.**
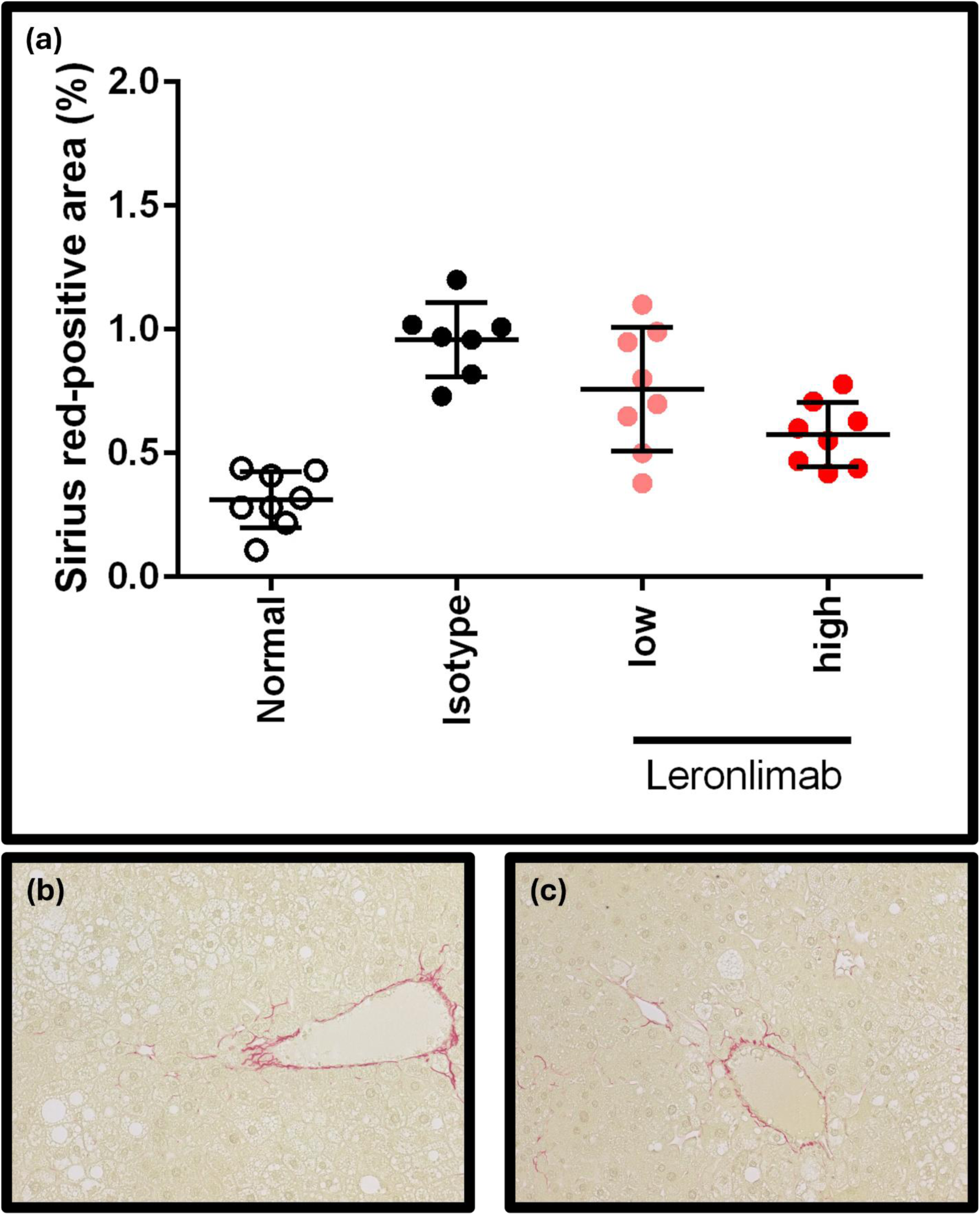
STAM model 1 fibrosis area and representative photomicrographs of Sirius red–stained liver sections. Panel a, Fibrosis area of the normal control versus isotype control versus leronlimab 5 and 10 mg/kg/week in the STAM model 1. P values by Bonferroni multiple comparison test: normal vs isotype control <0.0001; leronlimab low (5 mg/kg/week) vs isotype control 0.0937; leronlimab high (10 mg/kg/week) vs isotype control 0.0005. P values by Student’s t-test (one sided): normal vs isotype control <0.0001; leronlimab low (5 mg/kg/week) vs isotype control 0.0441; leronlimab high (10 mg/kg/week) vs isotype control <0.0001. Panel b, representative photomicrographs of Sirius red stained liver section for isotype control in the STAM model 1. Panel c, representative photomicrographs of Sirius red stained liver section for leronlimab 10 mg/kg in the STAM model 1.

### STAM model 2

As shown in Table 4 and 5, STAM model 2 was consistent with STAM model 1, with the isotype control group exhibiting significantly increased mean liver-to-body weight ratio compared with the normal group, with no significant differences observed between the isotype control group and the leronlimab-treated group. During the treatment period, one mouse died and two mice were euthanized in the isotype control group; no premature deaths or euthanasia occurred in the leronlimab-treated group.

**Table 4.** STAM model 2: between group differences in mouse parameters.

| **Comparison** | **Bonferroni Multiple**<br>**Comparison test (p-value)** | **Student's t-Test (p-**<br>**value)** |
| --- | --- | --- |
| **Liver-to-body weight ratio** |  |  |
| Normal vs isotype control | <0.0001 | <0.0001 |
| Leronlimab high vs isotype control | 0.2548 | 0.1056 |
| **Whole blood glucose** |  |  |
| Normal vs isotype control | <0.0001 | <0.0001 |
| Leronlimab high vs isotype control | 0.4156 | 0.1519 |
| **Plasma ALT** |  |  |
| Normal vs isotype control | 0.0568 | <0.0001 |
| Leronlimab high vs isotype control | 0.0127 | 0.0151 |
| **Plasma AST** |  |  |
| Normal vs isotype control | 0.9352 | 0.1473 |
| Leronlimab high vs isotype control | 0.0765 | 0.0281 |
| **Plasma triglyceride** |  |  |
| Normal vs isotype control | 0.0007 | <0.0001 |
| Leronlimab high vs isotype control | 0.9349 | 0.2823 |
| Leronlimab high = 10 mg/kg/week. |  |  |

**Table 5.**
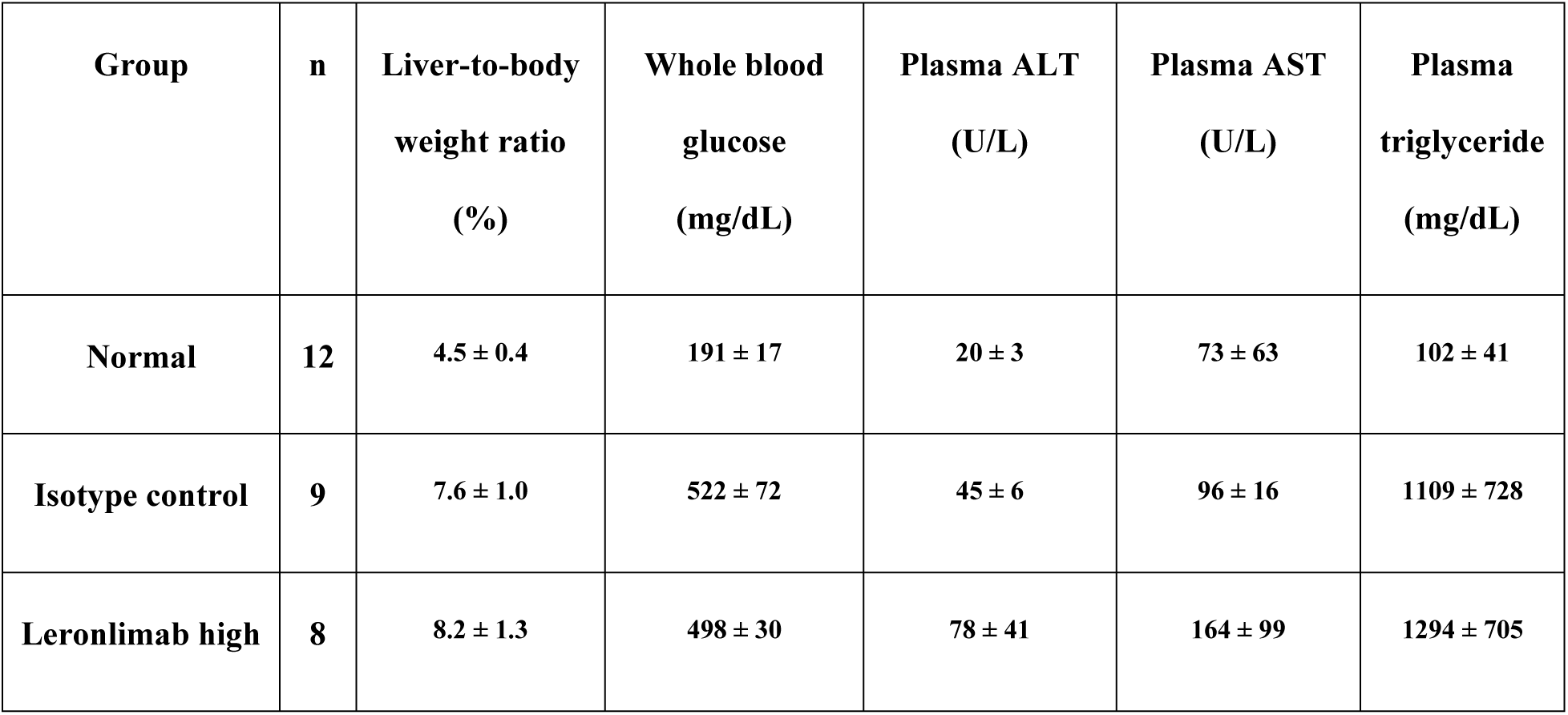
STAM model 2: Mean ± SD of each parameter.

Plasma alanine aminotransferase (ALT) and aspartate aminotransferase (AST) levels, whole blood glucose, and plasma triglyceride levels were significantly increased in the isotype control group compared with the normal group (Table 4 and 5). No significant differences were observed between the isotype control and leronlimab-treated groups for these biochemical parameters (Table 4 and 5).

As shown in Table 6, the NAS was significantly increased in the isotype control group compared with the normal group. Treatment with leronlimab resulted in a significant reduction in total NAS compared with the isotype control group, driven primarily by a significant reduction in hepatocyte ballooning.

**Table 6.** NAFLD activity score STAM model 2.

| Group | n | Score |  |  |  |  |  |  |  |  |  |  | NAS |
| --- | --- | --- | --- | --- | --- | --- | --- | --- | --- | --- | --- | --- | --- |
|  |  |  |  |  |  |  |  |  |  |  |  |  | (Mean ± SD) |
|  |  | Steatosis |  |  |  | Lobular inflammation |  |  |  | Hepatocyte ballooning |  |  |  |
|  |  | 0 | 1 | 2 | 3 | 0 | 1 | 2 | 3 | 0 | 1 | 2 |  |
| Normal | 12 | 12 | - | - | - | 12 | - | - | - | 12 | - | - | 0.0 ± 0.0 |
| Isotype control | 9 | 1 | 8 | - | - | - | - | 5 | 4 | 1 | 4 | 4 | 4.7 ± 0.5 |
| Leronlimab high | 12 | 6 | 5 | 1 | - | - | - | 5 | 7 | 11 | 1 | - | 3.3 ± 0.9 |

Leronlimab high = 10 mg/kg/week.
**Steatosis** – P values by Bonferroni multiple comparison test: normal vs isotype control 0.0002; leronlimab high (10 mg/kg/week) vs isotype control 0.2514. P values by Student's t-test: normal vs isotype control <0.0001; leronlimab high (10 mg/kg/week) vs isotype control 0.1126.
**Lobular inflammation** – P values by Bonferroni multiple comparison test: normal vs isotype control <0.0001; leronlimab high (10 mg/kg/week) vs isotype control 0.9052. P values by Student's t-test: normal vs isotype control <0.0001; leronlimab high (10 mg/kg/week) vs isotype control 0.2760.
**Hepatocyte ballooning** – P values by Bonferroni multiple comparison test: normal vs isotype control <0.0001; leronlimab high (10 mg/kg/week) vs isotype control <0.0001. P values by Student's t-test: normal vs isotype control <0.0001; leronlimab high (10 mg/kg/week) vs isotype control <0.0001.

Histological assessment by Sirius red staining demonstrated a significant reduction in fibrosis area in the leronlimab-treated group compared with the isotype control group (Fig. 2a).

**Fig. 2.**
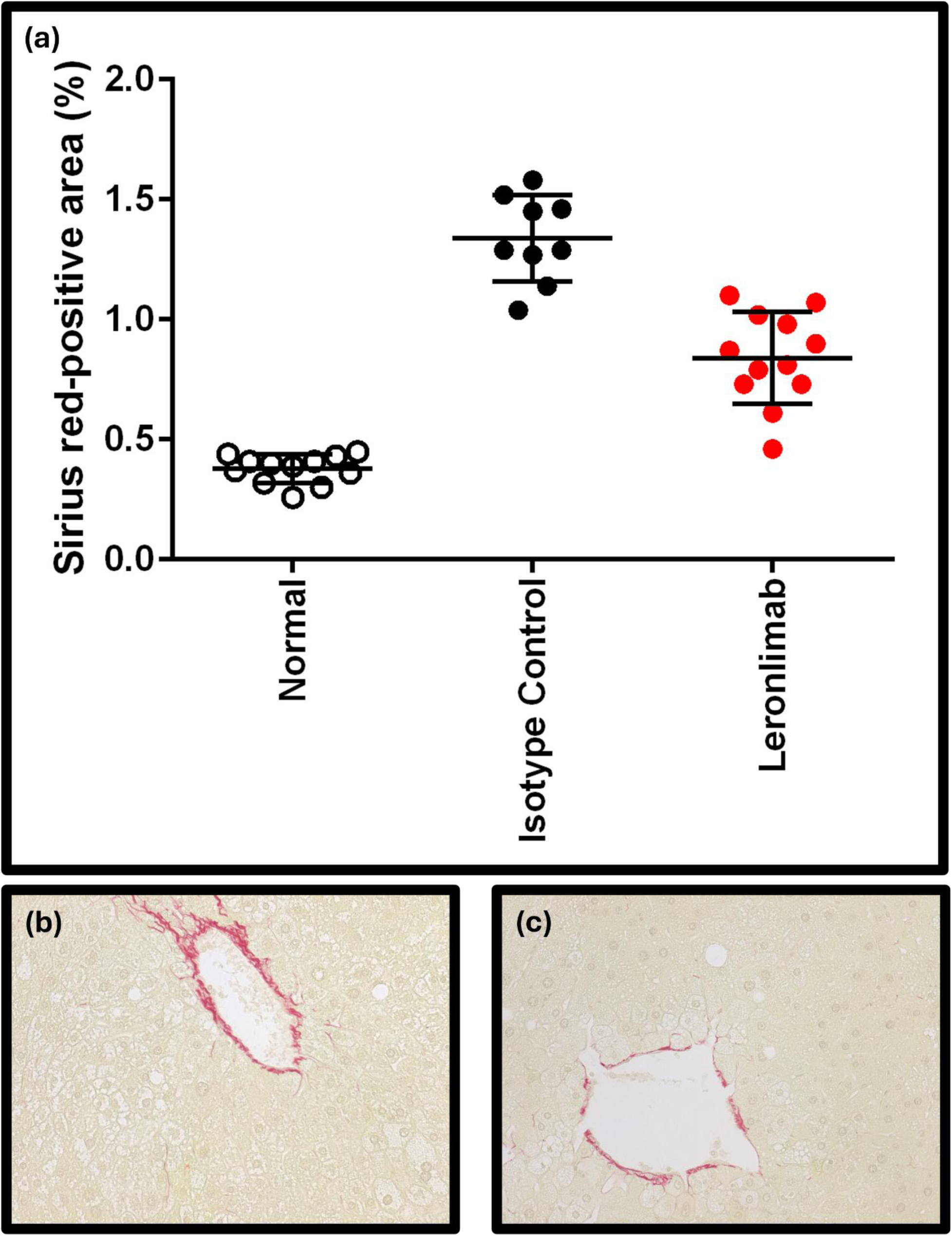
STAM model 2 fibrosis area and representative photomicrographs of Sirius red–stained liver sections. Panel a, Fibrosis area of the normal control versus isotype control versus leronlimab 10 mg/kg/week in the STAM model 2. P values by Bonferroni multiple comparison test: normal vs isotype control <0.0001; leronlimab high (10 mg/kg/week) vs isotype control <0.0001. P values by Student’s t-test (one-sided): normal vs isotype control <0.0001; leronlimab high (10 mg/kg/week) vs isotype control <0.0001. Panel b, representative photomicrographs of Sirius red stained liver section for isotype control in the STAM model 2. Panel c, representative photomicrographs of Sirius red stained liver section for leronlimab 10 mg/kg in the STAM model 2.

Representative Sirius red–stained liver sections for the isotype control and leronlimab (10 mg/kg/week) groups are shown in Fig. 2b and 2c, respectively. Compared with the normal group, increased fibrosis area was observed in both treatment groups.

### CCl₄-induced liver fibrosis model

As shown in Table 7 and 8, results from the CCl₄-induced liver fibrosis model were consistent with those observed in the STAM models. The isotype control group exhibited a significant increase in mean liver-to-body weight ratio compared with the normal group. No significant differences in liver-to-body weight ratio were observed between the isotype control and leronlimab-treated groups. No premature deaths or euthanasia occurred during the study.

**Table 7.** CCl4 model: between group differences in parameters.

| Comparison | Bonferroni Multiple<br>Comparison test (p-value) | Student's t-Test (p-value) |
| --- | --- | --- |
| **Liver-to-body weight ratio** |  |  |
| Normal vs isotype control | 0.0569 | 0.0132 |
| Leronlimab high vs isotype control | >0.9999 | 0.3666 |
| **Plasma ALT** |  |  |
| Normal vs isotype control | <0.0001 | <0.0001 |
| Leronlimab high vs isotype control | >0.9999 | 0.3484 |
| Leronlimab high = 10 mg/kg/week. |  |  |

**Table 8.** CCl4 model: Mean ± SD of each parameter.

| Group | n | Liver-to-body<br>weight ratio<br>(%) | Plasma ALT<br>(U/L) |
| --- | --- | --- | --- |
| Normal | 12 | 4.4 ± 0.3 | 14 ± 2 |
| Isotype control | 12 | 4.7 ± 0.4 | 25 ± 6 |
| Leronlimab high | 12 | 4.8 ± 0.4 | 26 ± 3 |

Plasma alanine aminotransferase (ALT) levels were significantly increased in the CCl₄-treated isotype control group compared with the non–CCl₄-treated normal group, consistent with CCl₄-induced hepatotoxicity. No significant differences in plasma ALT levels were observed between the isotype control and leronlimab-treated groups (Table 7 and 8).

Hepatic hydroxyproline content did not differ significantly between the isotype control and leronlimab-treated groups by either Bonferroni multiple comparison testing or Student’s t-test (Supplementary Table S9).

In contrast, histological assessment of fibrosis by Sirius red staining demonstrated a significant increase in fibrosis area in the isotype control group compared with the normal group, and a significant reduction in fibrosis area in the leronlimab-treated group compared with the isotype control group (Fig. 3a). Representative Sirius red–stained liver sections from the isotype control and leronlimab-treated (10 mg/kg/week) groups are shown in Fig. 3b and 3c, respectively. Compared with the normal group, increased fibrosis area was observed in both CCl₄-exposed groups.

**Fig. 3.**
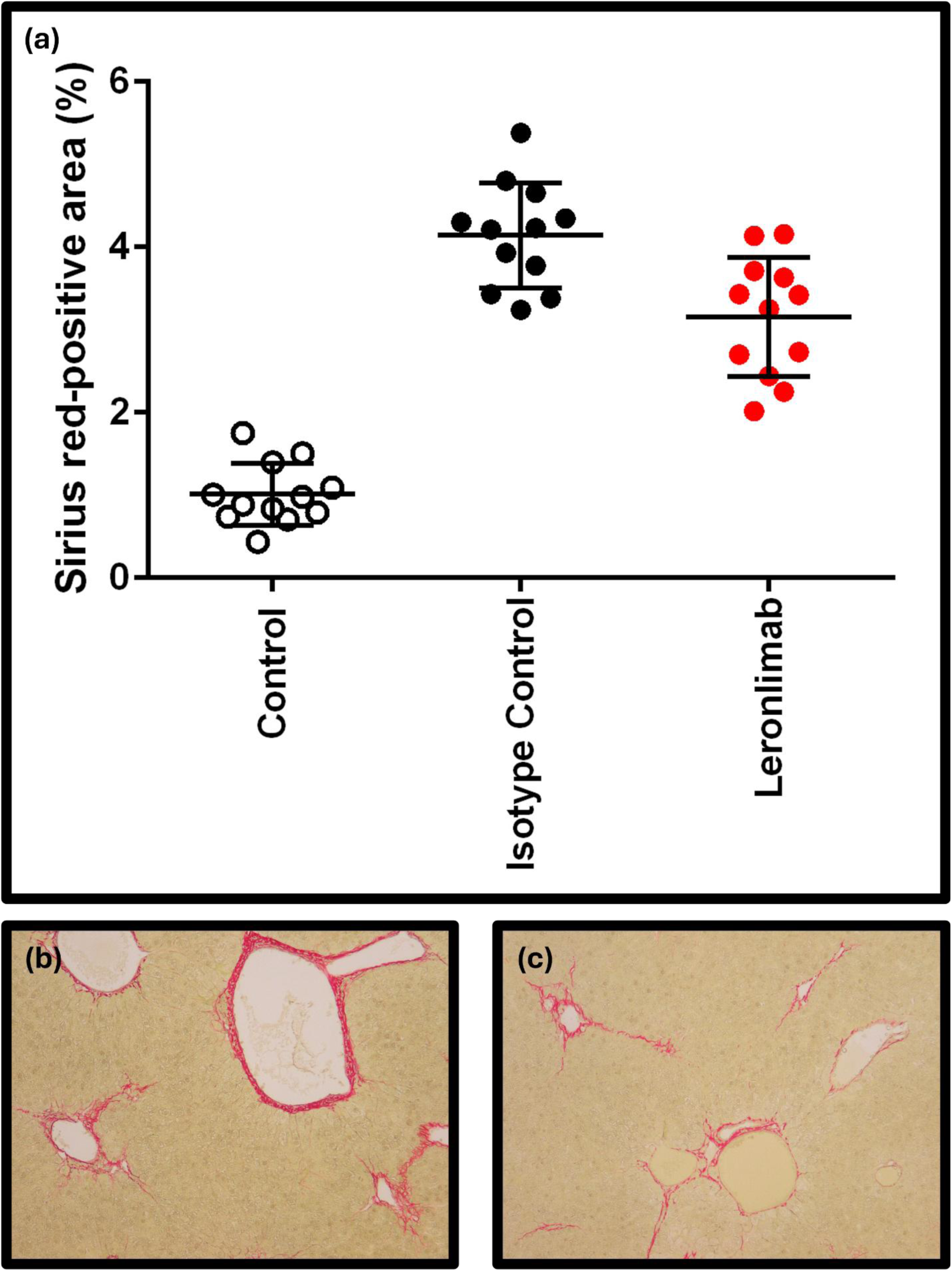
CCl₄-induced liver fibrosis model: fibrosis area and representative photomicrographs of Sirius red–stained liver sections. Panel a, fibrosis area of the normal control group versus isotype control versus leronlimab 10 mg/kg/week in the CCl_4_ mouse model. P values by Bonferroni multiple comparison test: normal vs isotype control <0.0001; leronlimab high (10 mg/kg/week) vs isotype control 0.0006. P values by Student’s t-test 9one-sided): normal vs isotype control <0.0001; leronlimab high (10 mg/kg/week) vs isotype control 0.0009. Panel b, representative photomicrographs of Sirius red-stained liver sections for isotype control in the CCl_4_ mouse model. Panel c, representative photomicrographs of Sirius red-stained liver sections for leronlimab 10 mg/kg in the CCl_4_ mouse model.

## Discussion

In the present study, leronlimab, a CCR5-blocking monoclonal antibody, was evaluated in two mechanistically distinct preclinical mouse models of liver fibrosis, including a metabolic injury–driven model (STAM) and a toxin-induced model (CCl₄). Across these models, leronlimab treatment was consistently associated with a significant reduction in histologically quantified fibrosis area, supporting a potential role for CCR5 blockade in modulating fibrogenic processes in the liver. To our knowledge, this study represents the first systematic evaluation of the antifibrotic effects of leronlimab in established preclinical models of liver fibrosis.

In both STAM model 1 and the confirmatory STAM model 2, treatment with leronlimab resulted in a significant reduction in fibrosis area compared with isotype control. Although total NAFLD activity score (NAS) was reduced in both STAM studies, improvements in individual NAS components were limited. In STAM model 1, reductions in NAS reflected combined numerical changes across steatosis, inflammation, and ballooning, whereas in STAM model 2 the reduction in NAS was driven primarily by a significant decrease in hepatocyte ballooning. Variability in the assessment of ballooning, which is known to exhibit substantial intra- and interobserver variability in both experimental models and human MASH, as well as differences in sample size between the two studies, may have contributed to these observations.

Notably, reductions in fibrosis area were observed in the absence of significant improvements in steatosis or inflammatory parameters, including serum transaminases. Given the established sequence of liver injury, inflammation, and fibrosis, these findings do not imply that fibrosis was modulated independently of inflammation. Rather, they suggest that CCR5 blockade by leronlimab may influence fibrogenic pathways involved in the maintenance or progression of established fibrosis, potentially through effects on immune cell recruitment, macrophage–stellate cell interactions, or extracellular matrix remodeling. These interpretations are consistent with prior studies implicating the CCL5–CCR5 axis in hepatic fibrogenesis via effects on macrophages and hepatic stellate cells.

The antifibrotic effects observed in the CCl₄-induced liver fibrosis model further support the consistency of leronlimab’s effects across distinct etiologies of liver injury. The CCl₄ model is characterized by hepatocellular injury, inflammatory cytokine release, and progressive fibrosis and is widely used to assess antifibrotic interventions. In this model, leronlimab significantly reduced fibrosis area by Sirius red staining, despite no significant changes in serum ALT levels, a finding that parallels the observations in the STAM models.

Several limitations of this study should be acknowledged. The STAM model develops mild to moderate perisinusoidal and pericellular fibrosis and does not progress to advanced bridging fibrosis. As such, the magnitude of antifibrotic effects observed here may not fully reflect potential effects in more advanced disease stages. Nevertheless, the fibrosis pattern observed in STAM mice resembles the perisinusoidal fibrosis characteristic of early to intermediate stages of human MASH, supporting the relevance of this model for evaluating antifibrotic interventions in this disease context.

In addition, while hydroxyproline quantification provides a measure of total hepatic collagen content, it does not capture spatial localization of fibrosis and may include collagen from non-pathological regions. This methodological limitation may explain the lack of significant changes in hydroxyproline content in the CCl₄ model despite clear reductions in fibrosis area assessed by Sirius red staining.

Importantly, this study did not directly assess cellular or molecular mechanisms underlying the observed antifibrotic effects, such as changes in hepatic stellate cell activation, macrophage infiltration, or extracellular matrix turnover. Future studies incorporating immunohistochemical and molecular analyses will be required to elucidate the precise mechanisms by which CCR5 blockade modulates fibrogenic pathways in the liver.

In conclusion, the present preclinical findings demonstrate that leronlimab treatment is associated with a significant reduction in liver fibrosis across multiple mouse models. These results support further investigation of leronlimab, either as monotherapy or in combination with other agents, for chronic liver diseases with fibrosis of varied etiologies.

### Data availability statement

CytoDyn Inc is committed to responsible data sharing regardless of the study outcome. Access to the data may be granted for legitimate requests as long as the data are not part of an ongoing or planned future regulatory submission. Requests for data and/or other study related information supporting this study should be made to CytoDyn at.

## CRediT author statement

Melissa Palmer: Writing – Original draft, Writing – Review and Editing

Taishi Hashiguchi: Methodology, Validation, Formal analysis, Investigation, Data curation, Writing – Review and Editing, Visualization, Supervision

A. Cyrus Arman: Conceptualization, Resources, Writing – Review and Editing, Supervision, Funding acquisition

Yuka Shirakata: Validation, Formal analysis, Investigation, Resources, Writing – Review and Editing, Visualization, Supervision

Jacob P. Lalezari: Conceptualization, Investigation, Writing – Review and Editing, Supervision, Funding acquisition

Neil E. Buss: Writing – Review and Editing

## Additional information

**Supplementary Information**. The online version contains supplementary material available at https:

## REFERENCES

1. Chan WK, Chuah KH, Rajaram RB, Lim LL, Ratnasingam J, Vethakkan SR. Metabolic Dysfunction-Associated Steatotic Liver Disease (MASLD): A State-of-the-Art Review. J Obes Metab Syndr. 2023;32(3):197–213; doi: 10.7570/jomes23052.

2. Lam BP, Bartholomew J, Bau S, Gilles H, Keller A, Moore A, et al. Focused Recommendations for the Management of Metabolic Dysfunction-Associated Steatohepatitis (MASH) by Advanced Practice Providers in the United States. J Clin Gastroenterol. 2025;59(4):298–309; doi: 10.1097/mcg.0000000000002140.

3. Tham EKJ, Tan DJH, Danpanichkul P, Ng CH, Syn N, Koh B, et al. The Global Burden of Cirrhosis and Other Chronic Liver Diseases in 2021. Liver Int. 2025;45(3):e70001; doi: 10.1111/liv.70001.

4. Younossi ZM, Kalligeros M, Henry L. Epidemiology of metabolic dysfunction-associated steatotic liver disease. Clin Mol Hepatol. 2025;31(Suppl):S32–s50; doi: 10.3350/cmh.2024.0431.

5. Younossi ZM, Stepanova M, Ong J, Trimble G, AlQahtani S, Younossi I, et al. Nonalcoholic Steatohepatitis Is the Most Rapidly Increasing Indication for Liver Transplantation in the United States. Clin Gastroenterol Hepatol. 2021;19(3):580–9.e5; doi: 10.1016/j.cgh.2020.05.064.

6. Younossi ZM, Wong G, Anstee QM, Henry L. The Global Burden of Liver Disease. Clin Gastroenterol Hepatol. 2023;21(8):1978–91; doi: 10.1016/j.cgh.2023.04.015.

7. Harrison SA, Bedossa P, Guy CD, Schattenberg JM, Loomba R, Taub R, et al. A Phase 3, Randomized, Controlled Trial of Resmetirom in NASH with Liver Fibrosis. N Engl J Med. 2024;390(6):497–509; doi: 10.1056/NEJMoa2309000.

8. Sanyal AJ, Newsome PN, Kliers I, Østergaard LH, Long MT, Kjær MS, et al. Phase 3 Trial of Semaglutide in Metabolic Dysfunction-Associated Steatohepatitis. N Engl J Med. 2025;392(21):2089–99; doi: 10.1056/NEJMoa2413258.

9. Campana L, Iredale JP. Regression of Liver Fibrosis. Semin Liver Dis. 2017;37(1):1–10; doi: 10.1055/s-0036-1597816.

10. Friedman SL, Rockey DC, McGuire RF, Maher JJ, Boyles JK, Yamasaki G. Isolated hepatic lipocytes and Kupffer cells from normal human liver: morphological and functional characteristics in primary culture. Hepatology. 1992;15(2):234–43; doi: 10.1002/hep.1840150211.

11. Barashi N, Weiss ID, Wald O, Wald H, Beider K, Abraham M, et al. Inflammation-induced hepatocellular carcinoma is dependent on CCR5 in mice. Hepatology. 2013;58(3):1021–30; doi: 10.1002/hep.26403.

12. Schwabe RF, Bataller R, Brenner DA. Human hepatic stellate cells express CCR5 and RANTES to induce proliferation and migration. Am J Physiol Gastrointest Liver Physiol. 2003;285(5):G949–58; doi: 10.1152/ajpgi.00215.2003.

13. Seki E, De Minicis S, Gwak GY, Kluwe J, Inokuchi S, Bursill CA, et al. CCR1 and CCR5 promote hepatic fibrosis in mice. J Clin Invest. 2009;119(7):1858–70; doi: 10.1172/jci37444.

14. Wen Y, Lambrecht J, Ju C, Tacke F. Hepatic macrophages in liver homeostasis and diseases-diversity, plasticity and therapeutic opportunities. Cell Mol Immunol. 2021;18(1):45–56; doi: 10.1038/s41423-020-00558-8.

15. Bertola A, Bonnafous S, Anty R, Patouraux S, Saint-Paul MC, Iannelli A, et al. Hepatic expression patterns of inflammatory and immune response genes associated with obesity and NASH in morbidly obese patients. PLoS One. 2010;5(10):e13577; doi: 10.1371/journal.pone.0013577.

16. Aydın MM, Akçalı KC. Liver fibrosis. Turk J Gastroenterol. 2018;29(1):14–21; doi: 10.5152/tjg.2018.17330.

17. Berres ML, Koenen RR, Rueland A, Zaldivar MM, Heinrichs D, Sahin H, et al. Antagonism of the chemokine CCL5 ameliorates experimental liver fibrosis in mice. J Clin Invest. 2010;120(11):4129–40; doi: 10.1172/jci41732.

18. Hammerich L, Tacke F. Hepatic inflammatory responses in liver fibrosis. Nat Rev Gastroenterol Hepatol. 2023;20(10):633–46; doi: 10.1038/s41575-023-00807-x.

19. Kumar S, Duan Q, Wu R, Harris EN, Su Q. Pathophysiological communication between hepatocytes and non-parenchymal cells in liver injury from NAFLD to liver fibrosis. Adv Drug Deliv Rev. 2021;176:113869; doi: 10.1016/j.addr.2021.113869.

20. Tong G, Chen X, Lee J, Fan J, Li S, Zhu K, et al. Fibroblast growth factor 18 attenuates liver fibrosis and HSCs activation via the SMO-LATS1-YAP pathway. Pharmacol Res. 2022;178:106139; doi: 10.1016/j.phrs.2022.106139.

21. Bradshaw D, Abramowicz I, Bremner S, Verma S, Gilleece Y, Kirk S, et al. Hepmarc: A 96 week randomised controlled feasibility trial of add-on maraviroc in people with HIV and non-alcoholic fatty liver disease. PLoS One. 2023;18(7):e0288598; doi: 10.1371/journal.pone.0288598.

22. Brata VD, Tacke F. Fatty liver disease: time to target CCR5? Expert Opin Ther Targets. 2024;28(5):335–9; doi: 10.1080/14728222.2024.2366880.

23. Friedman SL, Ratziu V, Harrison SA, Abdelmalek MF, Aithal GP, Caballeria J, et al. A randomized, placebo-controlled trial of cenicriviroc for treatment of nonalcoholic steatohepatitis with fibrosis. Hepatology. 2018;67(5):1754–67; doi: 10.1002/hep.29477.

24. Anstee QM, Neuschwander-Tetri BA, Wai-Sun Wong V, Abdelmalek MF, Rodriguez-Araujo G, Landgren H, et al. Cenicriviroc Lacked Efficacy to Treat Liver Fibrosis in Nonalcoholic Steatohepatitis: AURORA Phase III Randomized Study. Clin Gastroenterol Hepatol. 2024;22(1):124–34.e1; doi: 10.1016/j.cgh.2023.04.003.

25. Palmer M, Arman AC, Lawitz EJ, Sanchez WE, Mittal S, Hassanein T, et al. A Proof-of-Concept Phase 2a Partly Randomised Study Evaluating Leronlimab in Patients With Presumed Non-Cirrhotic Metabolic Dysfunction–Associated Steatohepatitis. Liver int commun. 2025;6:e70028; doi: 10.1002/lci2.70028.

26. Food and Agriculture Organization of the United Nartions. FAOLEX Database. Accessed June 25, 2025. Available from: https://www.fao.org/faolex/results/details/en/c/LEX-FAOC158111/. (2025). Accessed.

27. The Japanese Association of Laboratory Animal Facilities of National University Corporations. Accessed June 25, 2025. Available from: https://www.kokudoukyou.org/pdf/kisoku/sisin_eng.pdf. (2025). Accessed.

28. Science Council of Japan. Guidelines for Proper Conduct of Animal Experiments. Accessed June 25, 2025. Available from: https://www.scj.go.jp/ja/info/kohyo/pdf/kohyo-20-k16-2e.pdf. (2025). Accessed.

29. Chang ML, Yeh CT, Chang PY, Chen JC. Comparison of murine cirrhosis models induced by hepatotoxin administration and common bile duct ligation. World J Gastroenterol. 2005;11(27):4167–72; doi: 10.3748/wjg.v11.i27.4167.

30. Weber LW, Boll M, Stampfl A. Hepatotoxicity and mechanism of action of haloalkanes: carbon tetrachloride as a toxicological model. Crit Rev Toxicol. 2003;33(2):105–36; doi: 10.1080/713611034.

31. Kleiner DE, Brunt EM, Van Natta M, Behling C, Contos MJ, Cummings OW, et al. Design and validation of a histological scoring system for nonalcoholic fatty liver disease. Hepatology. 2005;41(6):1313–21; doi: 10.1002/hep.20701.

