## Supplemental materials for "Leronlimab a humanized anti-CCR5 monoclonal antibody ameliorates hepatic fibrosis in two preclinical fibrosis mouse models"

This appendix has been provided by the authors to give readers additional information about their work.

**Fig. S1. STAM models 1 and 2 and $CCl_4$-induced fibrosis model trial designs.**

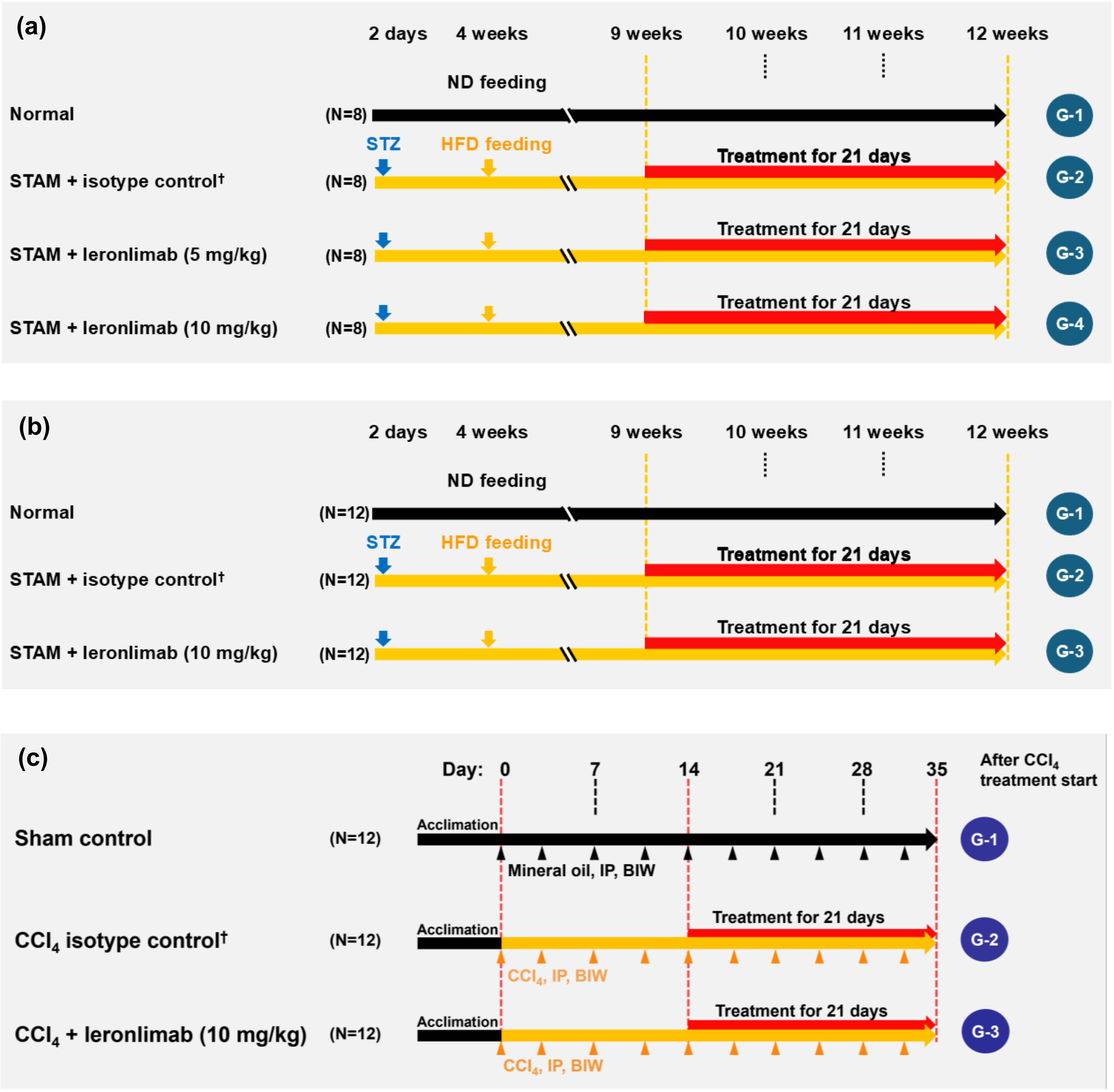


Legend for Fig. 1 – Schematic design of the three animal models. Panel A, STAM models 1 trial design. Panel B, STAM models 2 trial design. Panel C, $CCl_4$-induced fibrosis model trial design. † Ultra-LEAF purified human IgG4 isotype control recombinant antibody. BIW = twice per week; $CCl_4$ = carbon tetrachloride; HFD = high fat diet; IP = intra peritoneal; ND = normal diet; STAM = stelic animal model; STZ = streptozotocin.

**Table S1. Test item, lot number and storage conditions.**

| Test item/vehicle | Lot number and storage conditions |
|---|---|
| Leronlimab final drug product (FDP), 175 mg/mL | Manufacturer: Samsung BioLogics Co., Ltd.<br><br>Material specifications number: SPEC-0008 (Rev. 01)<br><br>Item/part number: PC2AA (SBL)<br><br>Lot number: 2100416<br><br>Storage: 2–8°C |
| Formulation buffer for dilution (placebo to match leronlimab) | Manufacturer: Samsung BioLogics Co., Ltd.<br><br>Material specifications number: SPEC-0009 (Rev. 00)<br><br>Item/part number: PC2B (SBL)<br><br>Lot number: 2100540<br><br>Storage: 2–8°C |
| Ultra-LEAF purified human IgG4 isotype control recombinant antibody (isotype control) | Manufacturer: BioLegend<br><br>Other names: IGHG4, immunoglobulin heavy chain constant region gamma 4<br><br>Isotype: human IgG4<br><br>Antibody type: recombinant<br><br>Host species: human<br><br>Formulation: 0.2 µm filtered in phosphate-buffered solution, pH 7.2, containing no preservative<br><br>Lot number: B425679<br><br>Storage: 2–8°C |

**Table S2. Definition of NAFLD activity score (NAS) components.**

| Item | Extent | Score |
|---|---|---|
| Steatosis | Steatosis at 50-fold magnification | |
| | <5% | 0 |
| | 5-33% | 1 |
| | >33-66% | 2 |
| | >66% | 3 |
| Lobular inflammation | Estimation of inflammatory | |
| | foci No foci | 0 |
| | <2 foci/200x | 1 |
| | 2-4 foci/200x | 2 |
| | >4 foci/200x | 3 |
| Ballooning | Estimation of number of ballooning cells | |
| | None | 0 |
| | Few ballooning cells | 1 |
| | Many cells/prominent ballooning | 2 |

**Supplemental Table S3. STAM model 1 individual data**

| Group | Mouse ID | Body weight (g) | Liver weight (mg) | Liver-to-body weight ratio (%) | Whole blood glucose (mg/dL) | Plasma ALT (U/L) | Plasma AST (U/L) | Plasma triglyceride (mg/dL) | Triglyceride (mg/g liver) | Steatosis | Inflammation | Ballooning | NAS | Sirius red-positive area (%) | Oil red-positive area (%) |
|---|---|---|---|---|---|---|---|---|---|---|---|---|---|---|---|
| Normal | 101 | 28.5 | 1416 | 5.0 | 159 | 19 | 53 | 89 | 4.0 | 0 | 0 | 0 | 0 | 0.32 | 3.49 |
| | 102 | 28.2 | 1326 | 4.7 | 183 | 18 | 39 | 73 | 5.9 | 0 | 0 | 0 | 0 | 0.41 | 1.32 |
| | 103 | 27.6 | 1257 | 4.6 | 207 | 24 | 43 | 88 | 7.5 | 0 | 0 | 0 | 0 | 0.22 | 1.60 |
| | 104 | 26.8 | 1290 | 4.8 | 192 | 19 | 46 | 172 | 5.3 | 0 | 0 | 0 | 0 | 0.28 | 2.48 |
| | 105 | 29.0 | 1403 | 4.8 | 181 | 20 | 35 | 165 | 4.6 | 0 | 0 | 0 | 0 | 0.11 | 1.29 |
| | 106 | 26.9 | 1129 | 4.2 | 213 | 20 | 73 | 124 | 4.9 | 0 | 0 | 0 | 0 | 0.43 | 2.15 |
| | 107 | 26.5 | 1192 | 4.5 | 240 | 19 | 34 | 78 | 1.6 | 0 | 0 | 0 | 0 | 0.44 | 1.32 |
| | 108 | 26.9 | 1200 | 4.5 | 175 | 16 | 39 | 129 | 5.8 | 0 | 0 | 0 | 0 | 0.28 | 2.81 |
| Isotype Control | 201 | 25.2 | 1815 | 7.2 | 522 | 26 | 75 | 116 | 18.3 | 1 | 2 | 2 | 5 | 0.82 | 28.38 |
| | 202 | | | | | | | | | | | | | | |
| | 203 | 12.7 | 1248 | 9.8 | 476 | 30 | 168 | 3195 | 24.3 | 1 | 3 | 0 | 4 | 1.01 | 19.20 |
| | 204 | 15.9 | 1299 | 8.2 | 775 | 126 | 254 | 3027 | 36.3 | 2 | 3 | 0 | 5 | 0.73 | 19.78 |
| | 205 | 23.8 | 1752 | 7.4 | 609 | 29 | 67 | 835 | 16.8 | 0 | 3 | 1 | 4 | 0.96 | 22.51 |
| | 206 | 17.8 | 1931 | 10.8 | 520 | 30 | 60 | 4559 | 26.3 | 2 | 2 | 0 | 4 | 1.20 | 23.95 |
| | 207 | 21.1 | 1679 | 8.0 | 701 | 26 | 73 | 1581 | 11.2 | 1 | 2 | 2 | 5 | 1.02 | 25.06 |
| | 208 | 21.0 | 1521 | 7.2 | 594 | 44 | 93 | 243 | 14.2 | 1 | 3 | 1 | 5 | 0.97 | 24.72 |
| Leronlimab low | 301 | 25.2 | 1768 | 7.0 | 549 | 35 | 87 | 573 | 32.2 | 0 | 2 | 0 | 2 | 1.10 | 26.12 |
| | 302 | 18.3 | 1755 | 9.6 | 426 | 24 | 57 | 1052 | 90.7 | 1 | 2 | 0 | 3 | 0.80 | 19.96 |
| | 303 | 24.0 | 1679 | 7.0 | 562 | 26 | 66 | 124 | 38.0 | 1 | 2 | 0 | 3 | 0.38 | 24.08 |
| | 304 | 21.6 | 1708 | 7.9 | 754 | 25 | 56 | 518 | 35.8 | 1 | 2 | 0 | 3 | 0.95 | 27.87 |
| | 305 | 21.7 | 1737 | 8.0 | 514 | 33 | 59 | 610 | 31.3 | 1 | 2 | 0 | 3 | 0.50 | 31.20 |
| | 306 | 22.1 | 1672 | 7.6 | 787 | 24 | 65 | 906 | 22.4 | 0 | 1 | 1 | 2 | 0.70 | 20.18 |
| | 307 | 22.4 | 1878 | 8.4 | 645 | 21 | 62 | 987 | 38.0 | 1 | 2 | 2 | 5 | 0.99 | 34.40 |
| | 308 | 17.5 | 1569 | 9.0 | >900 | 41 | 159 | 452 | 33.1 | 1 | 3 | 0 | 4 | 0.65 | 41.80 |
| Leronlimab high | 401 | 23.2 | 1921 | 8.3 | 526 | 26 | 55 | 3317 | 31.7 | 1 | 1 | 0 | 2 | 0.63 | 21.29 |
| | 402 | 24.9 | 1674 | 6.7 | 682 | 30 | 73 | 62 | 57.3 | 0 | 1 | 1 | 2 | 0.60 | 35.06 |
| | 403 | 13.9 | 1486 | 10.7 | 414 | 38 | 135 | 5982 | 75.1 | 1 | 3 | 0 | 4 | 0.42 | 16.59 |
| | 404 | 22.4 | 1577 | 7.0 | 562 | 29 | 62 | 955 | 33.7 | 0 | 2 | 0 | 2 | 0.78 | 32.66 |
| | 405 | 25.3 | 1956 | 7.7 | 549 | 46 | 99 | 482 | 58.4 | 0 | 3 | 0 | 3 | 0.44 | 31.76 |
| | 406 | 21.0 | 1638 | 7.8 | 595 | 24 | 51 | 601 | 18.9 | 1 | 2 | 2 | 5 | 0.55 | 20.59 |
| | 407 | 22.6 | 1795 | 7.9 | 479 | 44 | 81 | 435 | 50.6 | 1 | 2 | 0 | 3 | 0.47 | 35.25 |
| | 408 | 20.9 | 1580 | 7.6 | 575 | 32 | 69 | 176 | 30.0 | 1 | 2 | 0 | 3 | 0.71 | 28.34 |

: dead

**Supplemental Table S4. STAM model 2 individual data**

| Group | Mouse ID | Body weight (g) | Liver weight (mg) | Liver-to-body weight ratio (%) | Whole blood glucose (mg/dL) | Plasma ALT (U/L) | Plasma AST (U/L) | Plasma triglyceride (mg/dL) | Triglyceride (mg/g liver) | Steatosis | Inflammation | Ballooning | NAS | Sirius red-positive area (%) | Oil red-positive area (%) |
|---|---|---|---|---|---|---|---|---|---|---|---|---|---|---|---|
| Normal | 101 | 24.8 | 1206 | 4.9 | 176 | 18 | 44 | 107 | 4.4 | 0 | 0 | 0 | 0 | 0.26 | 0.07 |
| | 102 | 24.2 | 1195 | 4.9 | 226 | 20 | 61 | 96 | 2.5 | 0 | 0 | 0 | 0 | 0.37 | 0.03 |
| | 103 | 23.7 | 1016 | 4.3 | 190 | 20 | 70 | 45 | 2.4 | 0 | 0 | 0 | 0 | 0.43 | 0.25 |
| | 104 | 24.2 | 1031 | 4.3 | 220 | 20 | 37 | 86 | 3.7 | 0 | 0 | 0 | 0 | 0.30 | 0.05 |
| | 105 | 25.0 | 1209 | 4.8 | 185 | 28 | 261 | 98 | 3.3 | 0 | 0 | 0 | 0 | 0.36 | 0.08 |
| | 106 | 24.0 | 1181 | 4.9 | 181 | 18 | 40 | 151 | 2.6 | 0 | 0 | 0 | 0 | 0.41 | 0.15 |
| | 107 | 24.8 | 1245 | 5.0 | 179 | 17 | 74 | 195 | 5.5 | 0 | 0 | 0 | 0 | 0.32 | 0.17 |
| | 108 | 25.2 | 1182 | 4.7 | 208 | 20 | 60 | 94 | 4.8 | 0 | 0 | 0 | 0 | 0.45 | 0.14 |
| | 109 | 24.8 | 1080 | 4.4 | 176 | 18 | 37 | 112 | 4.8 | 0 | 0 | 0 | 0 | 0.41 | 0.08 |
| | 110 | 25.1 | 1087 | 4.3 | 179 | 19 | 46 | 75 | 8.3 | 0 | 0 | 0 | 0 | 0.44 | 0.27 |
| | 111 | 23.3 | 811 | 3.5 | 182 | 21 | 102 | 47 | 6.2 | 0 | 0 | 0 | 0 | 0.40 | 0.21 |
| | 112 | 24.1 | 1066 | 4.4 | 189 | 19 | 38 | 115 | 2.8 | 0 | 0 | 0 | 0 | 0.39 | 0.06 |
| Isotype Control | 201 | 25.0 | 1846 | 7.4 | 585 | 44 | 107 | 439 | 32.7 | 1 | 2 | 2 | 5 | 1.04 | 30.44 |
| | 202 | 24.7 | 2013 | 8.1 | 559 | 57 | 116 | 1489 | 42.0 | 1 | 2 | 1 | 4 | 1.14 | 17.65 |
| | 203 | | | | | | | | | | | | | | |
| | 204 | | | | | | | | | | | | | | |
| | 205 | 25.6 | 1563 | 6.1 | 545 | 46 | 80 | 2033 | 27.1 | 0 | 2 | 2 | 4 | 1.29 | 13.64 |
| | 206 | 21.5 | 1391 | 6.5 | 636 | 35 | 73 | 147 | 21.6 | 1 | 2 | 2 | 5 | 1.45 | 13.31 |
| | 207 | | | | | | | | | | | | | | |
| | 208 | 19.4 | 1592 | 8.2 | 456 | 47 | 117 | 1388 | 38.1 | 1 | 3 | 1 | 5 | 1.52 | 25.87 |
| | 209 | 25.7 | 1949 | 7.6 | 412 | 47 | 99 | 1240 | 46.4 | 1 | 3 | 1 | 5 | 1.27 | 31.31 |
| | 210 | 20.7 | 1953 | 9.4 | 446 | 43 | 87 | 2000 | 32.0 | 1 | 3 | 0 | 4 | 1.29 | 26.84 |
| | 211 | 23.4 | 1605 | 6.9 | 543 | 50 | 103 | 121 | 42.4 | 1 | 2 | 2 | 5 | 1.46 | 11.46 |
| | 212 | 18.5 | 1434 | 7.8 | 520 | 40 | 79 | 1126 | 37.8 | 1 | 3 | 1 | 5 | 1.58 | 23.42 |
| Leronlimab | 301 | 24.0 | 1922 | 8.0 | 474 | 92 | 204 | 1276 | 30.7 | 1 | 2 | 0 | 3 | 0.73 | 26.74 |
| | 302 | 21.7 | 1550 | 7.1 | 487 | 74 | 127 | 576 | 38.9 | 0 | 2 | 0 | 2 | 0.90 | 37.02 |
| | 303 | 22.3 | 1606 | 7.2 | 499 | 57 | 110 | 1122 | 33.6 | 0 | 3 | 0 | 3 | 0.46 | 17.52 |
| | 304 | 21.3 | 2370 | 11.1 | 578 | 171 | 363 | 2297 | 52.7 | 2 | 3 | 0 | 5 | 1.02 | 9.36 |
| | 305 | 23.5 | 1919 | 8.2 | 486 | 44 | 74 | 959 | 38.4 | 1 | 2 | 0 | 3 | 0.81 | 36.91 |
| | 306 | 24.1 | 1546 | 6.4 | 508 | 145 | 366 | 683 | 38.2 | 0 | 3 | 0 | 3 | 1.07 | 20.57 |
| | 307 | 21.7 | 2004 | 9.2 | 501 | 89 | 157 | 222 | 41.1 | 0 | 2 | 0 | 2 | 0.79 | 17.66 |
| | 308 | 20.7 | 1927 | 9.3 | 498 | 53 | 103 | 2162 | 38.3 | 1 | 3 | 0 | 4 | 0.87 | 17.53 |
| | 309 | 23.9 | 1976 | 8.3 | 497 | 64 | 119 | 2413 | 32.7 | 1 | 3 | 0 | 4 | 1.10 | 24.82 |
| | 310 | 23.6 | 1791 | 7.6 | 506 | 44 | 100 | 1693 | 23.4 | 0 | 2 | 1 | 3 | 0.61 | 17.54 |
| | 311 | 23.2 | 1675 | 7.2 | 447 | 64 | 120 | 993 | 38.1 | 0 | 3 | 0 | 3 | 0.73 | 24.20 |
| | 312 | 19.2 | 1740 | 9.1 | 497 | 40 | 124 | 1132 | 45.1 | 1 | 3 | 0 | 4 | 0.98 | 33.72 |

(shaded): euthanasia

(diagonal line): dead

**Supplemental Table S5. CCl4 model individual data**

| Group | Mouse ID | Body weight (g) | Liver weight (mg) | Liver-to-body weight ratio (%) | Plasma ALT (U/L) | Plasma ALT (U/L) | Hydroxyproline (µg/mg total protein) | Sirius red-positive area (%) |
|---|---|---|---|---|---|---|---|---|
| Control | 101 | 23.1 | 1034 | 4.5 | 14 | 1.58 | 1.58 | 0.74 |
| | 102 | 23.2 | 989 | 4.3 | 17 | 1.11 | 1.11 | 1.40 |
| | 103 | 23.2 | 1108 | 4.8 | 12 | 1.76 | 1.76 | 1.01 |
| | 104 | 20.0 | 926 | 4.6 | 18 | 2.66 | 2.66 | 0.44 |
| | 105 | 21.6 | 971 | 4.5 | 12 | 2.58 | 2.58 | 0.89 |
| | 106 | 20.6 | 867 | 4.2 | 15 | 3.06 | 3.06 | 0.71 |
| | 107 | 23.1 | 913 | 4.0 | 15 | 2.28 | 2.28 | 1.09 |
| | 108 | 21.1 | 945 | 4.5 | 15 | 2.50 | 2.50 | 0.98 |
| | 109 | 22.9 | 1039 | 4.5 | 13 | 2.42 | 2.42 | 0.79 |
| | 110 | 21.6 | 828 | 3.8 | 13 | 2.31 | 2.31 | 1.51 |
| | 111 | 23.0 | 1065 | 4.6 | 16 | 3.00 | 3.00 | 0.84 |
| | 112 | 19.9 | 869 | 4.4 | 12 | 1.75 | 1.75 | 1.75 |
| Isotype Control | 201 | 21.8 | 1121 | 5.1 | 28 | 3.38 | 3.38 | 4.21 |
| | 202 | 21.8 | 1009 | 4.6 | 28 | 2.65 | 2.65 | 3.43 |
| | 203 | 21.8 | 908 | 4.2 | 22 | 1.72 | 1.72 | 5.38 |
| | 204 | 20.4 | 986 | 4.8 | 33 | 2.61 | 2.61 | 4.23 |
| | 205 | 21.1 | 1055 | 5.0 | 35 | 2.33 | 2.33 | 3.38 |
| | 206 | 18.6 | 862 | 4.6 | 27 | 3.51 | 3.51 | 3.24 |
| | 207 | 21.6 | 1111 | 5.1 | 21 | 2.86 | 2.86 | 4.66 |
| | 208 | 22.7 | 1079 | 4.8 | 29 | 2.70 | 2.70 | 3.78 |
| | 209 | 21.8 | 1120 | 5.1 | 20 | 2.57 | 2.57 | 4.35 |
| | 210 | 24.6 | 918 | 3.7 | 16 | 2.36 | 2.36 | 3.93 |
| | 211 | 20.5 | 1046 | 5.1 | 21 | 3.36 | 3.36 | 4.30 |
| | 212 | 21.3 | 1018 | 4.8 | 22 | 2.64 | 2.64 | 4.80 |
| Leronlimab | 301 | 21.7 | 1047 | 4.8 | 29 | 2.11 | 2.11 | 3.71 |
| | 302 | 20.6 | 1092 | 5.3 | 30 | 2.85 | 2.85 | 3.43 |
| | 303 | 20.7 | 1029 | 5.0 | 30 | 3.20 | 3.20 | 4.16 |
| | 304 | 20.0 | 780 | 3.9 | 26 | 2.68 | 2.68 | 3.42 |
| | 305 | 21.6 | 1077 | 5.0 | 25 | 2.72 | 2.72 | 2.44 |
| | 306 | 18.6 | 874 | 4.7 | 28 | 2.11 | 2.11 | 4.14 |
| | 307 | 20.9 | 1075 | 5.1 | 24 | 3.03 | 3.03 | 3.25 |
| | 308 | 22.0 | 1041 | 4.7 | 24 | 2.70 | 2.70 | 3.63 |
| | 309 | 21.2 | 918 | 4.3 | 29 | 3.49 | 3.49 | 2.25 |
| | 310 | 21.5 | 1028 | 4.8 | 24 | 1.81 | 1.81 | 2.73 |
| | 311 | 21.1 | 982 | 4.7 | 22 | 2.34 | 2.34 | 2.70 |
| | 312 | 21.7 | 1143 | 5.3 | 20 | 2.16 | 2.16 | 2.02 |

Supplemental Table S6. STAM model 1 effect side and 95% confidence interval

| **Body weight** | Normal | Isotype Control | Leronlimab low | Leronlimab high |
|---|---|---|---|---|
| Lower 95% CI of mean | 26.78 | 15.55 | 19.43 | 18.80 |
| Upper 95% CI of mean | 28.32 | 23.74 | 23.77 | 24.75 |
| Effect size (Cohen's d) (vs Isotype Control) | -2.61 | - | -0.56 | -0.55 |

| **Liver weight** | Normal | Isotype Control | Leronlimab low | Leronlimab high |
|---|---|---|---|---|
| Lower 95% CI of mean | 1191 | 1366 | 1646 | 1561 |
| Upper 95% CI of mean | 1362 | 1847 | 1795 | 1846 |
| Effect size (Cohen's d) (vs Isotype Control) | 1.71 | - | -0.61 | -0.45 |

| **Liver-to-body weight ratio** | Normal | Isotype Control | Leronlimab low | Leronlimab high |
|---|---|---|---|---|
| Lower 95% CI of mean | 4.433 | 7.075 | 7.298 | 6.945 |
| Upper 95% CI of mean | 4.842 | 9.668 | 8.827 | 8.980 |
| Effect size (Cohen's d) (vs Isotype Control) | 3.95 | - | 0.26 | 0.31 |

| **Whole blood glucose** | Normal | Isotype Control | Leronlimab low | Leronlimab high |
|---|---|---|---|---|
| Lower 95% CI of mean | 172.5 | 500.5 | 508.7 | 481.2 |
| Upper 95% CI of mean | 215.0 | 698.6 | 775.5 | 614.3 |
| Effect size (Cohen's d) (vs Isotype Control) | 5.42 | - | -0.30 | 0.56 |

| **Plasma ALT** | Normal | Isotype Control | Leronlimab low | Leronlimab high |
|---|---|---|---|---|
| Lower 95% CI of mean | 17.48 | 10.68 | 22.85 | 26.79 |
| Upper 95% CI of mean | 21.27 | 78.17 | 34.40 | 40.46 |
| Effect size (Cohen's d) (vs Isotype Control) | 1.02 | - | 0.60 | 0.40 |

| **Plasma AST** | Normal | Isotype Control | Leronlimab low | Leronlimab high |
|---|---|---|---|---|
| Lower 95% CI of mean | 34.56 | 46.09 | 47.28 | 55.08 |
| Upper 95% CI of mean | 55.94 | 179.60 | 105.50 | 101.20 |
| Effect size (Cohen's d) (vs Isotype Control) | 1.36 | - | 0.67 | 0.66 |

| **Plasma triglyceride** | Normal | Isotype Control | Leronlimab low | Leronlimab high |
|---|---|---|---|---|
| Lower 95% CI of mean | 82.27 | 372.00 | 391.90 | -243.50 |
| Upper 95% CI of mean | 147.20 | 3501.00 | 913.60 | 3246.00 |
| Effect size (Cohen's d) (vs Isotype Control) | 1.58 | - | 1.10 | 0.23 |

| **Liver triglyceride** | Normal | Isotype Control | Leronlimab low | Leronlimab high |
|---|---|---|---|---|
| Lower 95% CI of mean | 3.519 | 13.130 | 22.620 | 28.740 |
| Upper 95% CI of mean | 6.381 | 28.980 | 57.750 | 60.190 |
| Effect size (Cohen's d) (vs Isotype Control) | 2.70 | - | -1.16 | -1.56 |

| **NAFLD Activity score** | Normal | Isotype Control | Leronlimab low | Leronlimab high |
|---|---|---|---|---|
| Lower 95% CI of mean | 0.0000 | 4.0770 | 2.2960 | 2.1060 |
| Upper 95% CI of mean | 0.0000 | 5.0660 | 3.9540 | 3.8940 |
| Effect size (Cohen's d) (vs Isotype Control) | 13.54 | - | 1.86 | 1.83 |

| **Steatosis score** | Normal | Isotype Control | Leronlimab low | Leronlimab high |
|---|---|---|---|---|
| Lower 95% CI of mean | 0.0000 | 0.5047 | 0.3630 | 0.1923 |
| Upper 95% CI of mean | 0.0000 | 1.7810 | 1.1370 | 1.0580 |
| Effect size (Cohen's d) (vs Isotype Control) | 2.31 | - | 0.50 | 0.83 |

| **Inflammation score** | Normal | Isotype Control | Leronlimab low | Leronlimab high |
|---|---|---|---|---|
| Lower 95% CI of mean | 0.000 | 2.077 | 1.553 | 1.368 |
| Upper 95% CI of mean | 0.000 | 3.066 | 2.447 | 2.632 |
| Effect size (Cohen's d) (vs Isotype Control) | 7.65 | - | 1.20 | 0.88 |

| **Ballooning score** | Normal | Isotype Control | Leronlimab low | Leronlimab high |
|---|---|---|---|---|
| Lower 95% CI of mean | 0.000 | 0.025 | -0.247 | -0.247 |
| Upper 95% CI of mean | 0.000 | 1.689 | 0.997 | 0.997 |
| Effect size (Cohen's d) (vs Isotype Control) | 1.47 | - | 0.63 | 0.63 |

| **Fibrosis area** | Normal | Isotype Control | Leronlimab low | Leronlimab high |
|---|---|---|---|---|
| Lower 95% CI of mean | 0.2158 | 0.8192 | 0.5506 | 0.4665 |
| Upper 95% CI of mean | 0.4067 | 1.0980 | 0.9669 | 0.6835 |
| Effect size (Cohen's d) (vs Isotype Control) | 5.00 | - | 0.95 | 2.72 |

| **Oil red-positive area** | Normal | Isotype Control | Leronlimab low | Leronlimab high |
|---|---|---|---|---|
| Lower 95% CI of mean | 1.373 | 20.420 | 22.060 | 21.630 |
| Upper 95% CI of mean | 2.742 | 26.320 | 34.590 | 33.750 |
| Effect size (Cohen's d) (vs Isotype Control) | 9.47 | - | -0.84 | -0.75 |

Supplemental Table S7. STAM model 2 effect side and 95% confidence interval

| **Body weight** | Normal | Isotype Control | Leronlimab |
|---|---|---|---|
| Lower 95% CI of mean | 24.05 | 20.60 | 21.45 |
| Upper 95% CI of mean | 24.82 | 24.85 | 23.42 |
| Effect size (Cohen's d) (vs Isotype Control) | -0.92 | - | 0.14 |

| **Liver weight** | Normal | Isotype Control | Leronlimab |
|---|---|---|---|
| Lower 95% CI of mean | 1032 | 1523 | 1686 |
| Upper 95% CI of mean | 1186 | 1888 | 1985 |
| Effect size (Cohen's d) (vs Isotype Control) | 3.32 | - | -0.55 |

| **Liver-to-body weight ratio** | Normal | Isotype Control | Leronlimab |
|---|---|---|---|
| Lower 95% CI of mean | 4.263 | 6.794 | 7.406 |
| Upper 95% CI of mean | 4.803 | 8.318 | 9.044 |
| Effect size (Cohen's d) (vs Isotype Control) | 4.20 | - | -0.57 |

| **Whole blood glucose** | Normal | Isotype Control | Leronlimab |
|---|---|---|---|
| Lower 95% CI of mean | 179.9 | 467.1 | 479.0 |
| Upper 95% CI of mean | 201.9 | 577.8 | 517.3 |
| Effect size (Cohen's d) (vs Isotype Control) | 6.82 | - | 0.46 |

| **Plasma ALT** | Normal | Isotype Control | Leronlimab |
|---|---|---|---|
| Lower 95% CI of mean | 18.04 | 40.69 | 51.90 |
| Upper 95% CI of mean | 21.63 | 50.20 | 104.30 |
| Effect size (Cohen's d) (vs Isotype Control) | 5.62 | - | -1.03 |

| **Plasma AST** | Normal | Isotype Control | Leronlimab |
|---|---|---|---|
| Lower 95% CI of mean | 32.78 | 83.00 | 101.00 |
| Upper 95% CI of mean | 112.20 | 108.30 | 226.80 |
| Effect size (Cohen's d) (vs Isotype Control) | 0.48 | - | -0.90 |

| **Plasma triglyceride** | Normal | Isotype Control | Leronlimab |
|---|---|---|---|
| Lower 95% CI of mean | 75.54 | 549.80 | 846.00 |
| Upper 95% CI of mean | 128.00 | 1669.00 | 1742.00 |
| Effect size (Cohen's d) (vs Isotype Control) | 2.13 | - | -0.26 |

| **Liver triglyceride** | Normal | Isotype Control | Leronlimab |
|---|---|---|---|
| Lower 95% CI of mean | 3.143 | 29.460 | 32.950 |
| Upper 95% CI of mean | 5.407 | 41.670 | 42.250 |
| Effect size (Cohen's d) (vs Isotype Control) | 5.87 | - | -0.27 |

| **NAFLD Activity score** | Normal | Isotype Control | Leronlimab |
|---|---|---|---|
| Lower 95% CI of mean | 0.000 | 4.282 | 2.700 |
| Upper 95% CI of mean | 0.000 | 5.051 | 3.800 |
| Effect size (Cohen's d) (vs Isotype Control) | 14.38 | - | 1.93 |

| **Steatosis score** | Normal | Isotype Control | Leronlimab |
|---|---|---|---|
| Lower 95% CI of mean | 0.0000 | 0.6327 | 0.1586 |
| Upper 95% CI of mean | 0.0000 | 1.1450 | 1.0080 |
| Effect size (Cohen's d) (vs Isotype Control) | 4.11 | - | 0.55 |

| **Inflammation score** | Normal | Isotype Control | Leronlimab |
|---|---|---|---|
| Lower 95% CI of mean | 0.0000 | 2.0390 | 2.2560 |
| Upper 95% CI of mean | 0.0000 | 2.8500 | 2.9110 |
| Effect size (Cohen's d) (vs Isotype Control) | 7.15 | - | -0.27 |

| **Ballooning score** | Normal | Isotype Control | Leronlimab |
|---|---|---|---|
| Lower 95% CI of mean | 0.0000 | 0.7898 | -0.1001 |
| Upper 95% CI of mean | 0.0000 | 1.8770 | 0.2667 |
| Effect size (Cohen's d) (vs Isotype Control) | 2.91 | - | 2.46 |

| **Fibrosis area** | Normal | Isotype Control | Leronlimab |
|---|---|---|---|
| Lower 95% CI of mean | 0.3409 | 1.2000 | 0.7177 |
| Upper 95% CI of mean | 0.4158 | 1.4750 | 0.9606 |
| Effect size (Cohen's d) (vs Isotype Control) | 7.71 | - | 2.68 |

| **Oil red-positive area** | Normal | Isotype Control | Leronlimab |
|---|---|---|---|
| Lower 95% CI of mean | 0.0785 | 15.6500 | 18.1200 |
| Upper 95% CI of mean | 0.1815 | 27.4500 | 29.1500 |
| Effect size (Cohen's d) (vs Isotype Control) | 4.30 | - | -0.25 |

Supplemental Table S8. $CCl_4$ model effect side and 95% confidence interval

| **Body weight** | Control | Isotype Control | Leronlimab |
|---|---|---|---|
| Lower 95% CI of mean | 21.12 | 20.60 | 20.37 |
| Upper 95% CI of mean | 22.77 | 22.40 | 21.56 |
| Effect size (Cohen's d) (vs Isotype Control) | -0.30 | - | 0.42 |

| **Liver weight** | Control | Isotype Control | Leronlimab |
|---|---|---|---|
| Lower 95% CI of mean | 907.50 | 964.20 | 941.70 |
| Upper 95% CI of mean | 1018.00 | 1075.00 | 1073.00 |
| Effect size (Cohen's d) (vs Isotype Control) | 0.64 | - | 0.13 |

| **Liver-to-body weight ratio** | Control | Isotype Control | Leronlimab |
|---|---|---|---|
| Lower 95% CI of mean | 4.215 | 4.470 | 4.546 |
| Upper 95% CI of mean | 4.568 | 5.013 | 5.054 |
| Effect size (Cohen's d) (vs Isotype Control) | 0.85 | - | -0.25 |

| **Plasma ALT** | Control | Isotype Control | Leronlimab |
|---|---|---|---|
| Lower 95% CI of mean | 13.05 | 21.54 | 23.83 |
| Upper 95% CI of mean | 15.61 | 28.79 | 28.01 |
| Effect size (Cohen's d) (vs Isotype Control) | 2.46 | - | -0.21 |

| **Liver hydroxyproline** | Control | Isotype Control | Leronlimab |
|---|---|---|---|
| Lower 95% CI of mean | 1.88 | 2.40 | 2.28 |
| Upper 95% CI of mean | 2.63 | 3.05 | 2.92 |
| Effect size (Cohen's d) (vs Isotype Control) | 0.85 | - | 0.24 |

| **Fibrosis area** | Normal | Isotype Control | Leronlimab |
|---|---|---|---|
| Lower 95% CI of mean | 0.7750 | 3.7390 | 2.7000 |
| Upper 95% CI of mean | 1.2500 | 4.5420 | 3.6130 |
| Effect size (Cohen's d) (vs Isotype Control) | 6.06 | - | 1.45 |

**Supplemental Table S9. Differences in liver hydroxyproline content between the isotype control group and leronlimab.**

| **Liver hydroxyproline (μg/mL)** | | **Bonferroni multiple comparison test (p value)** | **Student's t-test (two tailed p value)** |
|---|---|---|---|
| Control vs. isotype control | 2.25 ± 0.59 | 0.1849 | 0.0465 |
| Leronlimab vs. isotype control | 2.6 ± 0.50 | >0.9999 | 0.5532 |
